# Downregulation of the central noradrenergic system by *Toxoplasma gondii* infection

**DOI:** 10.1101/345900

**Authors:** Isra Alsaady, Ellen Tedford, Mohammad Alsaad, Greg Bristow, Shivali Kohli, Matthew Murray, Matthew Reeves, M.S. Vijayabaskar, Steven J. Clapcote, Jonathan Wastling, Glenn A. McConkey

## Abstract

The parasitic protozoan *Toxoplasma gondii* becomes encysted in brain and muscle tissue during chronic infection, a stage that was previously thought to be dormant but has been found to be active and associated with physiological effects in the host. Dysregulation of catecholamines in the CNS has previously been observed in chronically-infected animals. In the study described here, the noradrenergic system was suppressed with decreased levels of norepinephrine in brains of infected animals and in infected neuronal cells in vitro. Expression of dopamine β-hydroxylase (DBH), essential for synthesis of norepinephrine from dopamine, was the most differentially-expressed gene in infections *in vitro* and was down-regulated in infected brain tissue, particularly in the prefrontal cortex and dorsal locus coeruleus/pons region. The down-regulated DBH expression in infected rat catecholaminergic and human neuronal cells corresponded with decreased norepinephrine and increased dopamine. As the DBH suppression was observed *in vitro*, this effect is not caused by neuroinflammation. Silencing of DBH expression was specific for *T. gondii* infection and was not observed with CMV infection. The noradrenergic-linked behaviors of sociability and arousal were altered in chronically-infected animals, with a high correlation between DBH expression and infection intensity. These findings together provide a plausible mechanism to explain prior discrepancies in changes to CNS neurotransmitters levels with infection. The suppression of norepinephrine synthesis observed here may, in part, explain behavioural effects of infection, associations with mental illness, and neurological consequences of infection such as the loss of coordination and motor impairments associated with human toxoplasmosis.

## Introduction

*T. gondii* infects warm-blooded animals and is characterised by a transient acute infection wherein vegetative tachyzoite forms rapidly replicate in tissues followed by a persistent chronic infection. Chronic stages of infection can persist for years and potentially the lifetime of the host with the bradyzoite-stage parasites encysted in cells within immunoprivileged tissues, including muscle, eyes, and brain. Several reports have published host behavioral changes with infection. A selective loss of aversion to feline urine and increased motor activity has been observed in rodents, specifically manipulating behavior that will enhance the probability of parasite transmission (1, 2).

Toxoplasmosis can be a severe disease in immunocompromised individuals and *in utero*. Infection can cause retinochoroiditis and congenital hydrocephalus and cerebral calcifications. *T. gondii* was recently ranked the second most important food-borne parasite in Europe and is classified as a Neglected Parasitic Infection (CDC, Atlanta) (3). It has also been linked by epidemiological studies to cognitive impairment and major mental illnesses. Severe cases are associated with psychoses, seizures and loss of coordination. Yet there are currently no available cures for infection. Sensorimotor defects, tremors and headshaking have also been observed in chronically-infected mice (4, 5). In the brain, encysted bradyzoite-stage parasites are restricted to neurons, and recent work has found that neurons are the primary target cell for *T. gondii* during CNS infection (6, 7). As the parasite encysts in neurons, this study investigated changes in gene expression during neural cell infection.

Early studies found changes in dopaminergic neurotransmission associated with infection, with high levels of dopamine (DA) in brain tissue cysts and abrogation of infection-induced behavior changes when animals were treated with dopamine antagonists, haloperidol and GBR-12909 (8–10). Perturbations in catecholaminergic signalling with chronic infection have been observed, with elevated DA metabolites in the cortex and decreased NE in the cortex and amygdala and loss of amphetamine-induced locomotor activity (11,12). There are discrepancies in observations of changes in dopamine levels in the brain with *T. gondii* infection (13–17). Increased levels of dopamine in infected cells have been found when catecholaminergic cells are maintained at a physiological pH (8, 18–20). *T. gondii* contains two paralogous genes encoding tyrosine/phenylalanine hydroxylase, TgAAAH1 and TgAAAH2, that were recently found to be involved in cyst development in the cat intestine (21, 22). The genes are expressed in bradyzoites but mutants with one of the paralogs deleted had no effect on DA levels and did not disrupt behavior changes with infection (23). The role of these parasite genes in brain DA levels is the subject of a separate study. Here, we examined noradrenergic neurotransmission and, through examination of gene expression changes, identified a biological mechanism that not only provides a possible resolution for published findings on DA, but also describes a mechanism whereby NE is suppressed during CNS infection.

## Results

### Norepinephrine regulation in the brain during *T. gondii* infection

The effect of chronic infection on CNS NE and DA was monitored by measuring levels in the brains of *T. gondii*-infected animals. The level of NE was significantly changed with infection (p=0.0019) with a 50±14% decrease in NE level in the brain (Figure 1A). Decreased NE in *T. gondii*-infected mice has been observed in other studies (11, 13). The suppression observed with infection (Figure 1A) is analogous to decreases in CNS NE levels observed with high affinity DBH inhibitors (24). High doses of disulfiram and nepicastat, that have been used clinically, reduce brain NE levels by 36-45% (25, 26). Although NE was reduced with infection, the rats displayed no obvious signs of pathology, as commonly observed with chronic *T. gondii* infections in rats (27). The median level of DA in the brains of infected rats was increased to double the uninfected level in this cohort, but this was not statistically significant (Figure 1B, p=0.12). These observations fit with other investigations, in which high DA levels were observed in cysts but brain tissue levels of DA were unchanged (15, 18, 28)

**Figure 1:**
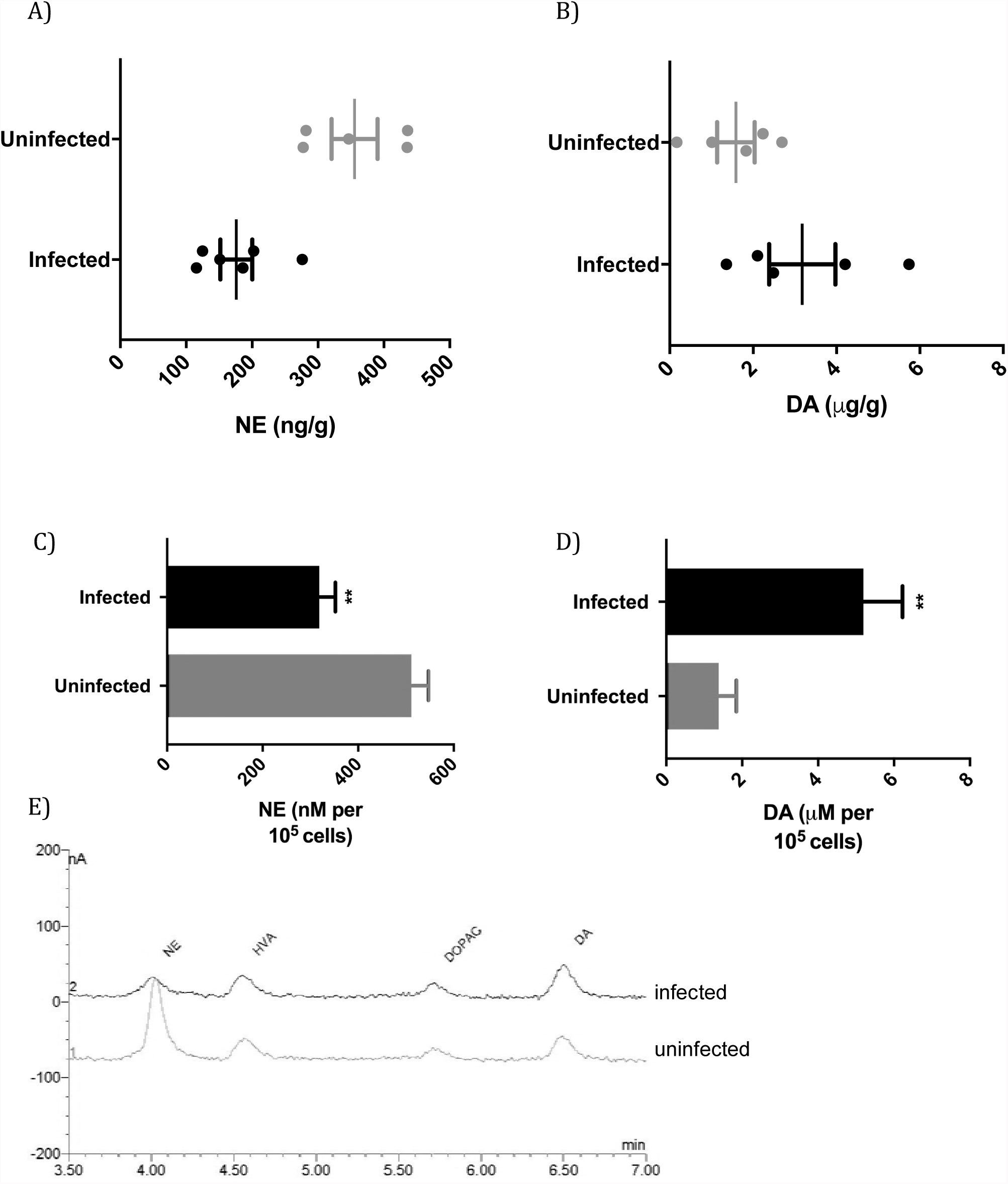
Infection effects on catecholamines in the brain and catecholaminergic cells. A) Norepinephrine levels in the brains of uninfected and *T. gondii*-infected rats (p=0.0019, Student’s t-test; n=11). B) Dopamine levels in the brains of uninfected and infected rats (p=0.12, Student’s t-test t; n=11). C) Norepinephrine levels of uninfected and infected catecholaminergic PC12 cells at day 5 of infection. Multiplicity of infection is 1; p=0.0024, n=3. D) Dopamine levels in infected PC12 cells plotted as above. p=0.0043. E) Chromatograms from HPLC-ED of uninfected and infected cells.

To determine whether the change in level of NE was a result of infection of neurons or was a consequence of infection such as due to the immune response, we performed infections with a model of catecholaminergic neurons, PC12 cells. PC12 cells synthesize and package the catecholamines DA, NE and, to a lesser extent, epinephrine for vesicle-mediated release and express dopamine receptors. To simulate chronic infection, we shocked the tachyzoites with high pH to induce bradyzoite development prior to infection of cells (8, 18). As DA synthesis by PC12 cells is sensitive to pH, this technique maintains the full DA capacity of the cells (20, 29).

NE and DA levels were measured in PC12 cells five days after parasite infection. NE levels were decreased in infected cultures to 62±6.1% (p=0.0024) of uninfected cell level (Figure 1C, 1E). The reduction in NE cannot be due to cell lysis as values are expressed relative to cell number. DA levels in infected PC12 cells were greater than uninfected cells (p=0.0043) in the same samples that exhibited suppression of NE (Figure 1D). The 3.8±0.74-fold increase is similar to that found in our previously published work with infected PC12 cells (8, 18). Hence, infection reduced NE whilst elevating dopamine levels.

Regulation of the levels of NE and DA may be due to changes in synthesis, transport and storage, or degradation. Further, the mechanism(s) responsible for the opposing decrease in NE and increase in DA in catecholaminergic cells was unclear from these observations. Therefore, we examined the effects of the parasite on proteins expressed by the host neuronal cells.

### Down-regulation of a key enzyme for norepinephrine synthesis during infection

To try to decipher the biological mechanism(s) responsible for the decreased NE in the brain with infection, a whole-genome transcriptome scan was performed. We used PC12 cells that were differentiated to form dendritic extensions and synapse- and neuronal-like functions to detect changes in expression of genes encoding neural proteins and processes (e.g. catecholamine synthesis and release, receptors). This permitted detection of neuronal genes which might not be possible in the mixture of cell types in infected brain samples (30). Further, in contrast to other transcriptome studies, parasites were shocked to induce bradyzoite development (18). Surprisingly, of the 26,405 rat genes detected, the most significantly altered expression was down-regulation of the dopamine β-hydroxylase (DBH) gene (p= 7.2×10^−13^, FDR= 2.3×10^−11^). Housekeeping gene expression (GAPDH, ribosomal proteins, tRNA ligases, tubulin) was unchanged in infected cells in the transcriptome screen, permitting detection of specific differentially expressed genes. Simultaneous analysis found up-regulated expression of *T. gondii* bradyzoite genes (BAG1, LDH2, MAG1, MIC13), although the number of reads was significantly lower than host cells. Prior transcriptomic studies of whole infected brain tissue have principally identified changes in expression of genes in the host immune response, as might be expected with the mixture of cell types in the brain (30, 31).

The effect in human neuronal cells of *T. gondii* infection on DBH gene expression was measured. The BE(2)-M17 cell line, derived from a human neuroblastoma and possessing catecholaminergic properties and neuritic processes, was infected and monitored over a five-day time course of infection, during which time chronic, bradyzoite stages of the parasite will develop. Expression of the DBH gene was downregulated 5.7±1.1-fold by day 3 of infection (p=0.00032) and 17±1.4-fold by day 5 of infection (p=0.0010) (Figure 2A). DBH levels were consistent in uninfected BE(2)-M17 cells throughout the experiment (one-way ANOVA, p=0.97). DBH down-regulation was also observed in BE(2)-M17 cells infected with the *T. gondii* ME49 strain (data not shown).

**Figure 2:**
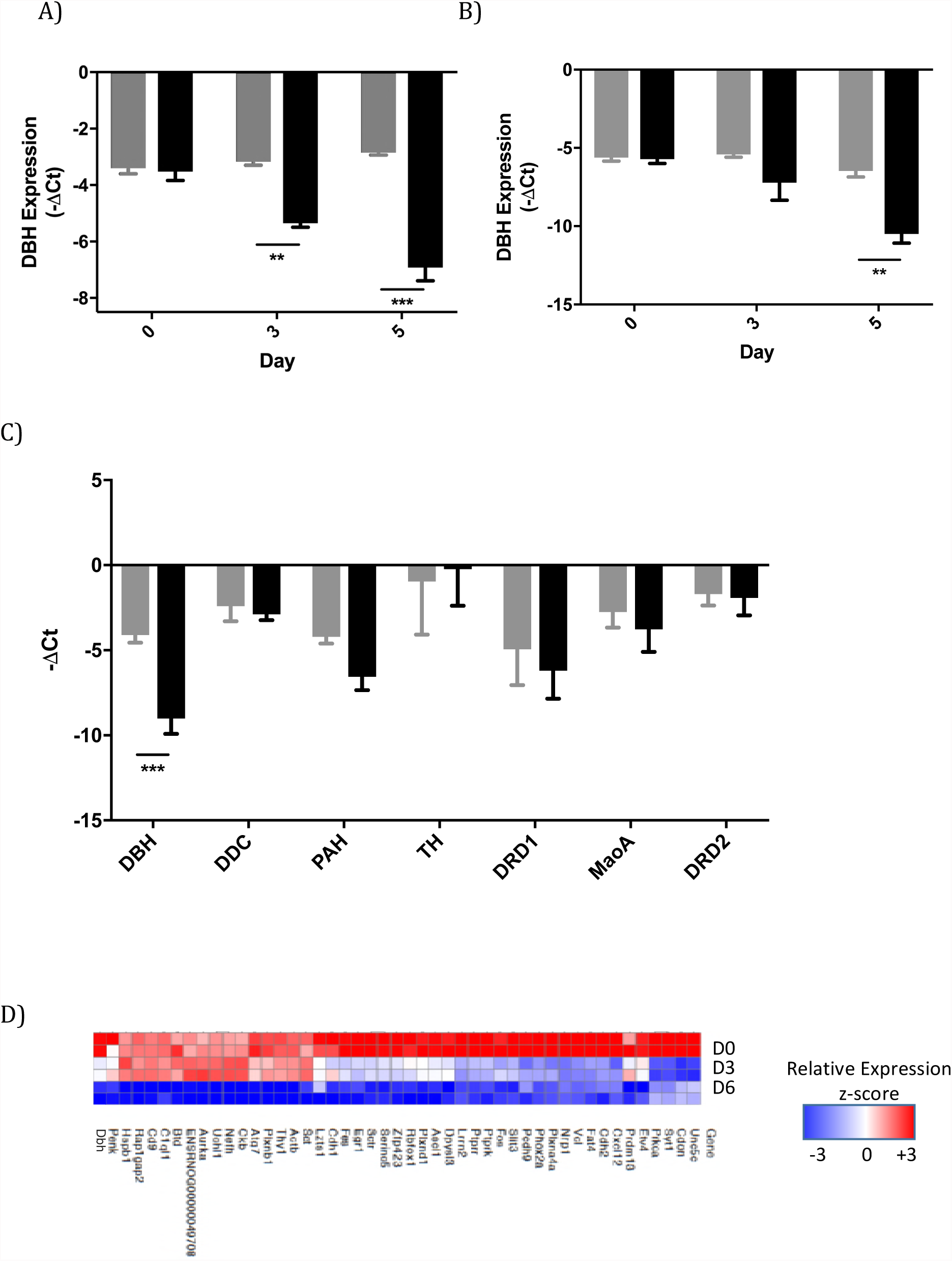
Norepinephrine biosynthesis in catecholaminergic cells with *T. gondii* infection. A) Dopamine ß-hydroxylase gene expression during infection (black) or control (grey) in human BE(2)-M17 neuronal cells over a time course of infection relative to GAPDH. Multiplicity of infection is 1; **, p=0.0010; ***, p=0.00032; n=3. B) Plot of the level of DBH mRNA over a time course of infection of rat catecholaminergic cells. **, p=0.0046, n=3. C) Expression of the set of catecholaminergic genes during infection (black) or uninfected (grey). Only the DBH gene expression was significantly altered by infection (n=3, multiple t tests, ***, p=0.008). DBH, dopamine β-hydroxylase; DDC, dopa decarboxylase; PAH, phenylalanine hydroxylase; TH, tyrosine hydroxylase; DRD1, dopamine receptor D1; MaoA, monoamine oxidase A; DRD2, dopamine receptor D2. Error bars are ±SEM. D) Heat map of down-regulated neurological gene expression in transcriptome analysis of infected cells at day 0, 3, and 6.

A time course of infection and DBH expression was repeated in PC12 cells. DBH gene expression was decreased after 72 hours of infection and further after 120 hours in PC12 cells (30±2-fold), relative to GAPDH (p=0.0046, n=3) (Figure 2B). Microscopic analysis verified the maintenance of cell numbers and viability during the time course experiments. The level of DBH mRNA in uninfected PC12 cells was unchanged over the course of the experiment (one-way ANOVA, p= 0.58).

We then surveyed a collection of catecholamine biosynthesis and metabolism genes for changes with infection. Quantitative analysis of gene expression in uninfected cells found that DBH down-regulation was the only significantly changed gene (Figure 2C). This concurs with the whole-genome RNA sequencing data. The decreased DBH gene expression was observed both in PC12 cell cultures and cultures with differentiated neuronal-like cells (with dendritic extensions that possess synaptic functions). Although expression of the phenylalanine hydroxylase gene (PAH) appeared reduced, this was not significant (p=0.06). Levels of mRNA for tyrosine hydroxylase, dopamine decarboxylase, monoamine oxidase A, and dopamine receptors D1 and D2 were unchanged with infection. The lack of change in rat tyrosine hydroxylase and dopamine decarboxylase gene expression with *T. gondii* infection corresponds with previously published data (8). In addition to DBH, a panel of genes were down-regulated >2-fold. These were enriched 2.0-2.9-fold for genes involved in neuron differentiation and development (Figure 2D, Table S1, p values 2.3-3.8 × 10^−6^). Some genes were up-regulated with infection, although at lower significance levels, with the most significant set involved in cellular stress responses related to immunity, as might be expected for an infectious agent (Table S1, p values 0.45-1.8 × 10^−6^).

DBH is the key link between NE and DA, with DBH metabolizing DA into NE. Decreased DBH will decrease synthesis of NE, and simultaneously increase levels of the precursor DA. Suppression of DBH by down-regulated expression of its gene provides a mechanistic explanation for the observed increase in DA in infected PC12 cells above (Figures 1C, 1D) coincident with decreased levels of NE. DA was not significantly increased in infected rat brains, as might be expected with the disproportionately smaller number of noradrenergic compared to dopaminergic neurons.

### Dopamine β-hydroxylase down-regulation suppresses norepinephrine in the brain

We examined whether the down-regulation of DBH gene expression in neuronal cells was detectable during *in vivo* infection. The level of DBH expression in the infected brain was examined. DBH mRNA was quantified in the brains of chronically-infected rats. Gene expression was down-regulated in infected animals by a median of 32±2.1-fold relative to uninfected animals (Figure 3A; p=0.0023). We examined the relationship between the intensity of brain infection and DBH expression. A strong negative correlation was observed in infected animals between DBH mRNA and cyst density (tissue cysts can contain thousands of bradyzoites), with a correlation coefficient of −0.90 (Table 1). The coefficient of determination (R^2^) of 0.82 is a good fit for the linear regression.

**Figure 3:**
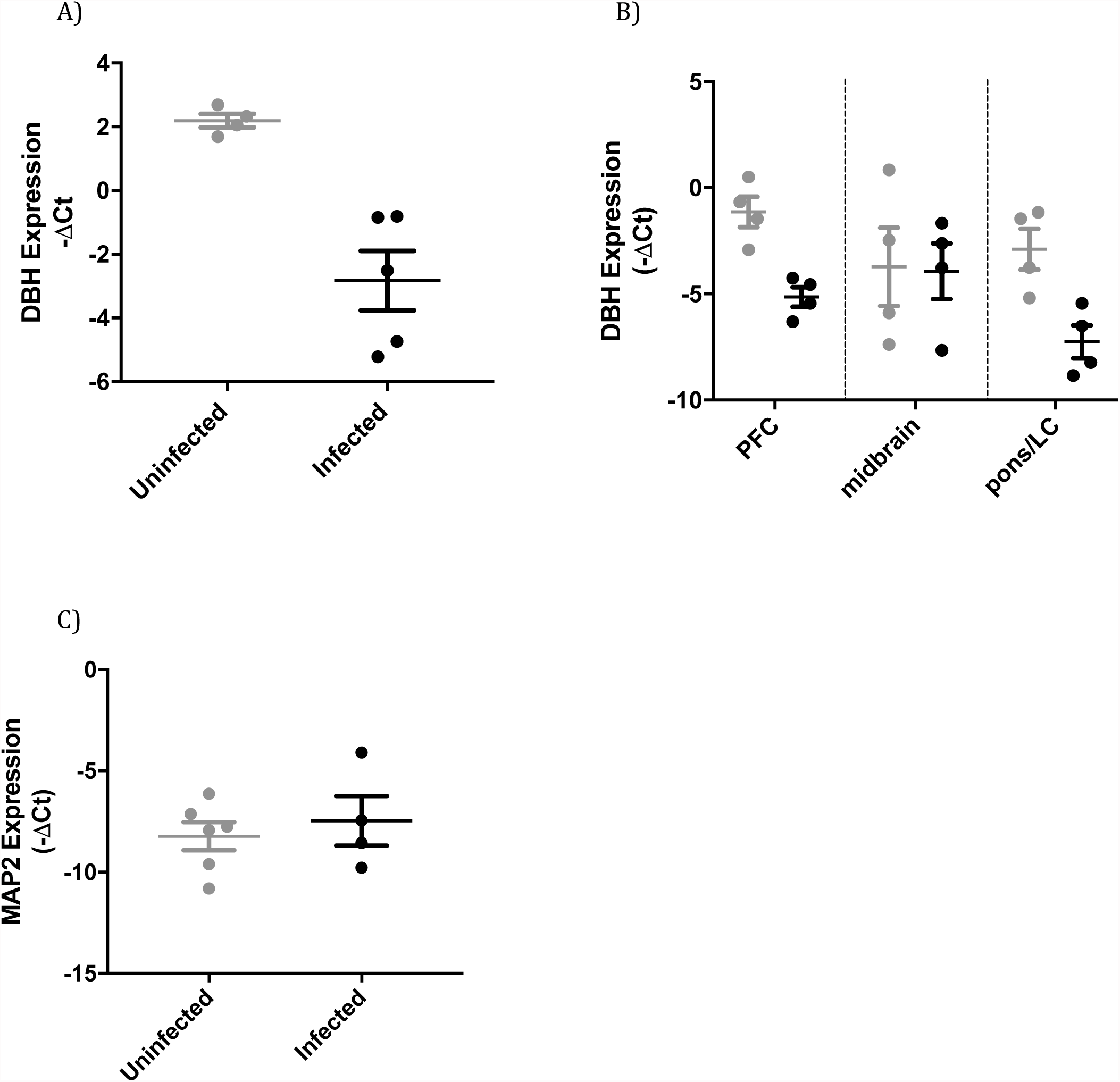
Infection downregulates dopamine ß-hydroxylase gene expression in the brain. A) DBH gene expression in the brains of uninfected (grey) and chronically-infected (black) male rats plotted relative to GAPDH (p=0.0023, student t test; n=9). B) Brain region specific DBH gene expression in uninfected and infected rats. PFC, prefrontal cortex; LC, locus coeruleus. Error bars are ±SEM. C) Plot showing expression of the neuronal MAP2 gene (as a percentage of GAPDH) in uninfected (grey) and chronically-infected (black) brains (p=0.57, Student’s t-test; n=10).

**Table 1.**
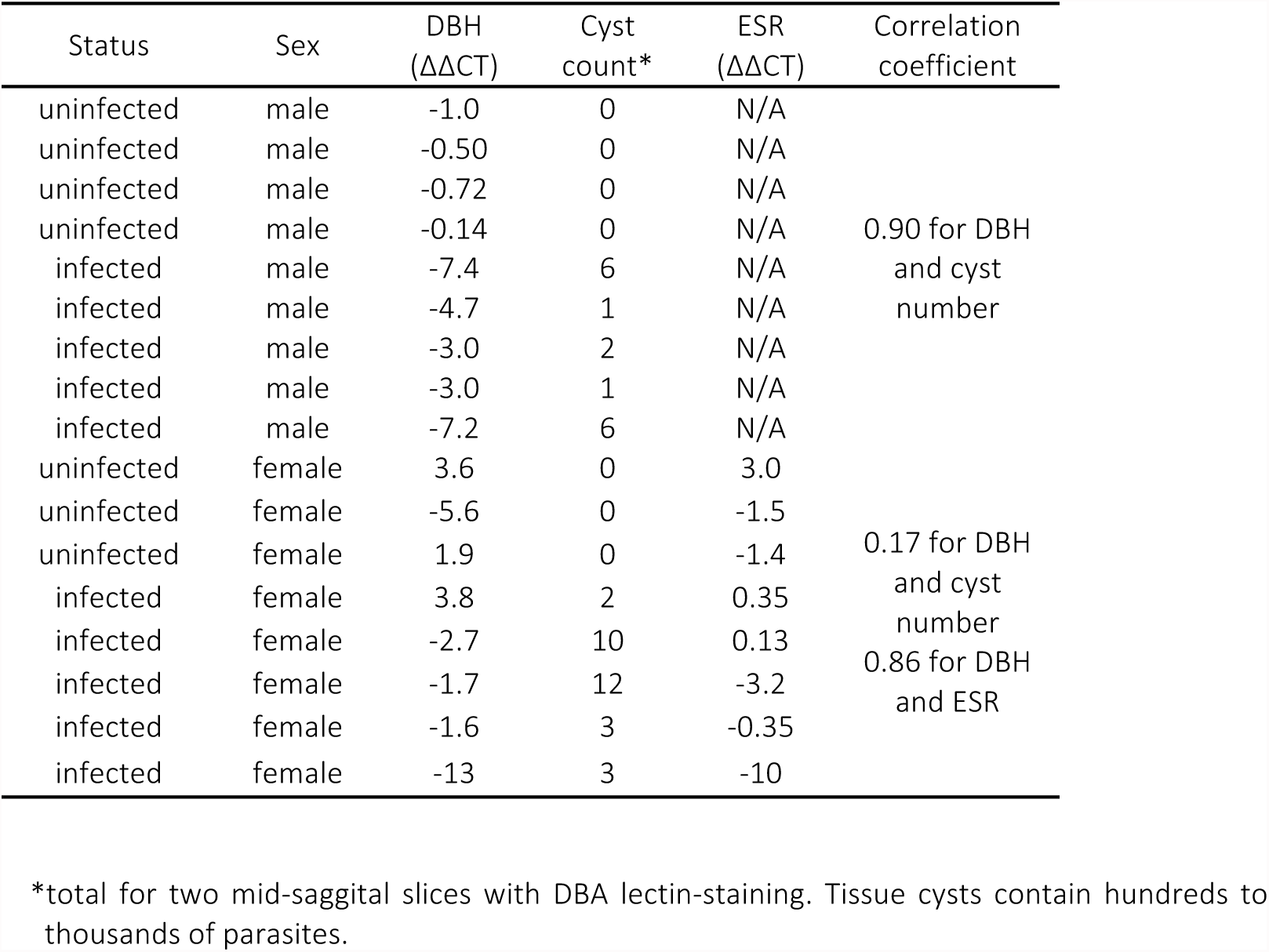
DBH and ESR gene expression in the CNS with chronic *T. gondii* infection

DBH is expressed in noradrenergic neurons in the CNS, principally in the locus coeruleus (LC) with efferents extending to most brain regions. Therefore, we examined DBH gene expression in different brain regions in infected animals. DBH mRNA levels were lower (p=0.0034 and 0.012, respectively) in the frontal lobe (prefrontal cortex (PFC)) and the dorsal region (containing the LC, cerebellum, pons, and surrounding tissue) in infected animals, whereas DBH expression was unchanged in the midbrain region containing the hippocampus, thalamus and hypothalamus (p=0.93) (Figure 3B). The posterior area and the PFC had 2.5-fold and 4.5-fold, respectively, lower DBH mRNA in infected rats.

One plausible alternative explanation for the decrease in NE in the infected rat brains could be poor neuronal health or neuronal death. *T. gondii* can lyse neurons and synaptic loss and neuronal dysfunction has been observed in infected mice (32). In this study, we found no difference in neurons between infected and uninfected rats based on quantification of a neuron-specific mRNA, that encoding microtubule-associated protein 2 (MAP2) (Figure 3C; p= 0.57).

### Suppressed dopamine β-hydroxylase alters norepinephrine-linked behaviors

A decrease in CNS NE, as observed with *T. gondii* infection (Figure 1A), may have specific effects on behavior. Sociability, arousal and anxiety are all behaviors associated with CNS noradrenergic signalling (33, 34). Rodents with NE deficiency exhibit increased sociability and lower arousal and anxiety levels. Cerebral NE levels were associated with social interest and male aggression (35). NE levels elevated by disruption of monoamine oxidase A result in increased aggression in mice (36). In contrast, aggressive behavior is decreased and social memory altered in *Dbh*−/− knockout mice (33). In this study, the three-chambered social approach test was used to measure sociability in uninfected and *T. gondii*-infected mice. This test is a well-established model for measuring social interactions in mouse models of autism (37).

In the first phase of the social approach test, which measures sociability, preference for exploring a cylinder containing a stranger mouse rather than an empty cylinder was measured (38). Chronically-infected mice explored the novel mouse for substantially longer times (median 31 s, range 3-91 s, n=27) than the uninfected mice (median 23 s, range 0.2-66 s, n=24), in line with lower NE levels (Figure 4A). The level of brain DBH mRNA in infected mice in the trials was significantly lower than the control mice (Figure 4B), particularly for the male mice (p=0.0032) where DBH was down-regulated 5.8±1.5-fold. The level of CNS DBH in the infected animals exhibited a negative correlation with the time of investigating the novel mouse, albeit a weak correlation (Figure S1). Infection has previously been associated with social interaction, with *T. gondii*-infected rats exhibiting a longer duration of social interaction than controls (39). The decreased DBH observed here provides an explanation for increased social interaction with *T. gondii* infection.

**Figure 4:**
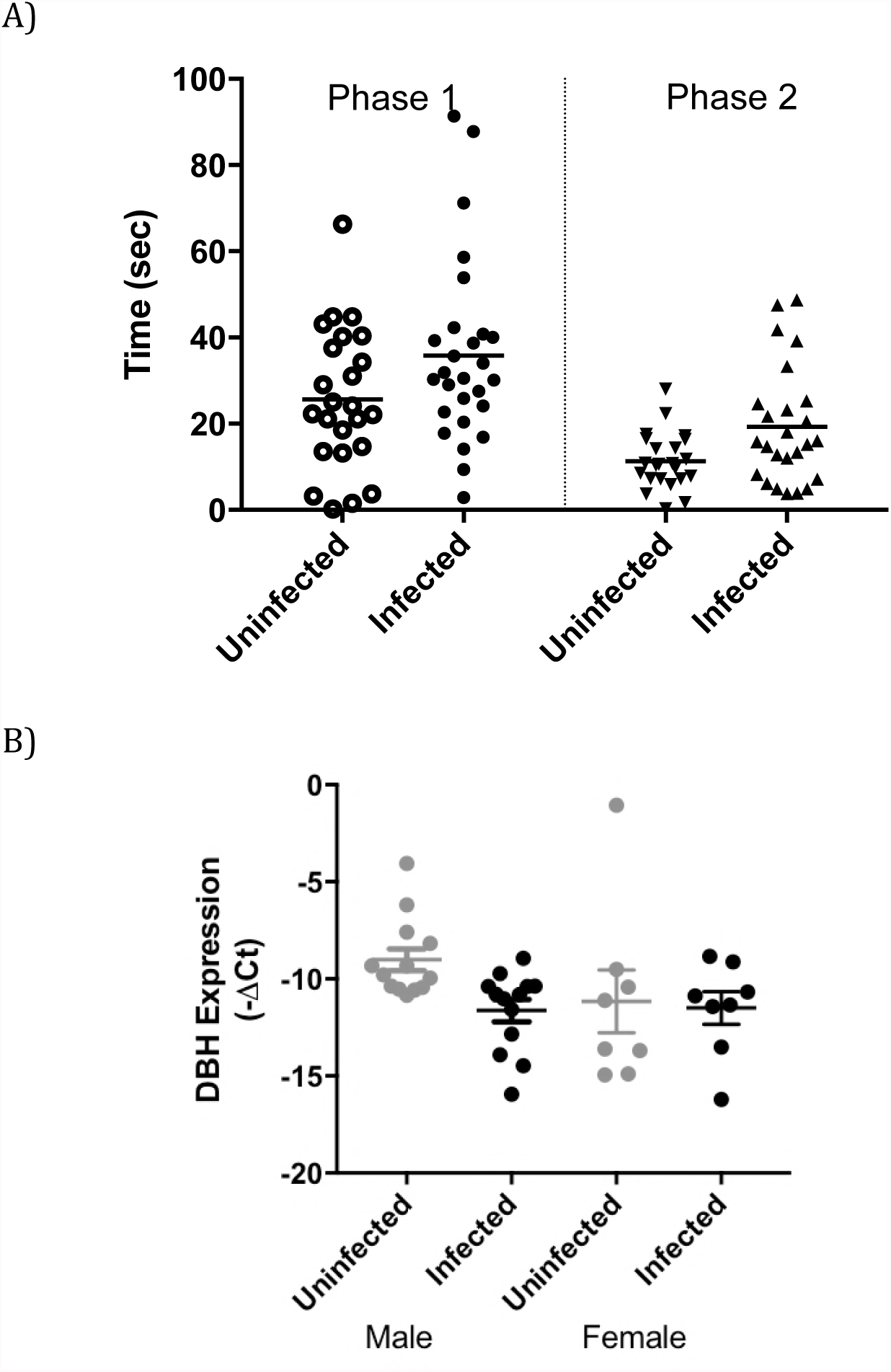
Social approach and dopamine ß-hydroxylase gene expression with *T. gondii* infection. A) A combined plot showing time spent (seconds) investigating a novel mouse (Stranger 1) in preference to an empty container in phase 1 of the test of uninfected (grey) and infected animals (black). In the second phase of the test, time spent investigating a second novel mouse (Stranger 2) in preference to the first stranger mouse was measured. For the two phases, the p values are 0.063 and 0.025, respectively. C) Expression of CNS DBH in the trial animals with male and female (p= 0.0032, n=26 and p=0.85, n=16, respectively). Error bars are ±SEM.

In Phase 2 of the social approach test, which measures preference for social novelty, mice encountered the Stranger 1 mouse (the now familiar mouse) as well as a novel mouse (Stranger 2) in the formerly empty cylinder. Although both uninfected and infected mice investigated the novel stranger, infected mice spent significantly longer in contact with the novel stranger, with medians of 15 s and 10.5 s for infected and control mice, respectively (Figure 4A; p=0.025). There was a wide range of values in these trials, with uninfected mice spending 0-28 s in contact with the novel mouse and infected mice spending 0.6-49 s. We examined the possibility of an association of DBH level in the infected mice with length of time investigating a novel mouse, but these parameters did not correlate.

Arousal is measured as a response to evoked or elicited activity and has been quantified in rodents by locomotion in a novel environment, such as an open field, at early time points (40). Locomotion of chronically-infected and uninfected mice in an open field apparatus was monitored and ambulation recorded over 1-min intervals to 5 minutes, then over 5-min intervals to 15 minutes. The mice were individually placed in the open field and allowed to settle for 60 seconds (minute 1), while the experimenter withdrew from the apparatus, before readings were taken. *T. gondii* infected mice exhibited decreased locomotor activity in the open field at early time points but not at later times (Figures 5A, S2). Uninfected mice travelled 3.1-3.2 m during minutes 2 and 3, whereas infected animals travelled 2.3-2.5 m. The differences in distances travelled were significant (p<0.0001 and 0.0015, respectively, for each reading). Representative tracking of uninfected and control mice illustrates the decreased locomotor activity during early time points (Figure 5B). The tracking in the figure also replicates the loss of fear of open spaces found in prior studies of *T. gondii*-infected rodents (41). After three minutes, infected and control groups showed similar levels of activity in the open field; p=0.91 and 0.27, respectively, for minutes 4 and 5. No decrease was observed between infected and uninfected mice in the 5-min intervals from minutes 5-15 (Figure S2), matching prior studies of locomotion in *T. gondii*-infected rodents monitored over longer periods (circa 30 minutes) (41–43). In previous studies, mobility during 1-minute intervals was not reported, and hence changes in initial behavioral response or arousal would not be observed. The DBH mRNA levels in the mice exhibited a moderate negative correlation with early locomotor activity (Figure S3), with a Pearson’s correlation coefficient of −0.48. Published studies of *Dbh*−/− knockout mice have described attenuated arousal and decreased locomotion, similar to that observed here, in ambulation in an open field at early time points (33, 34).

**Figure 5:**
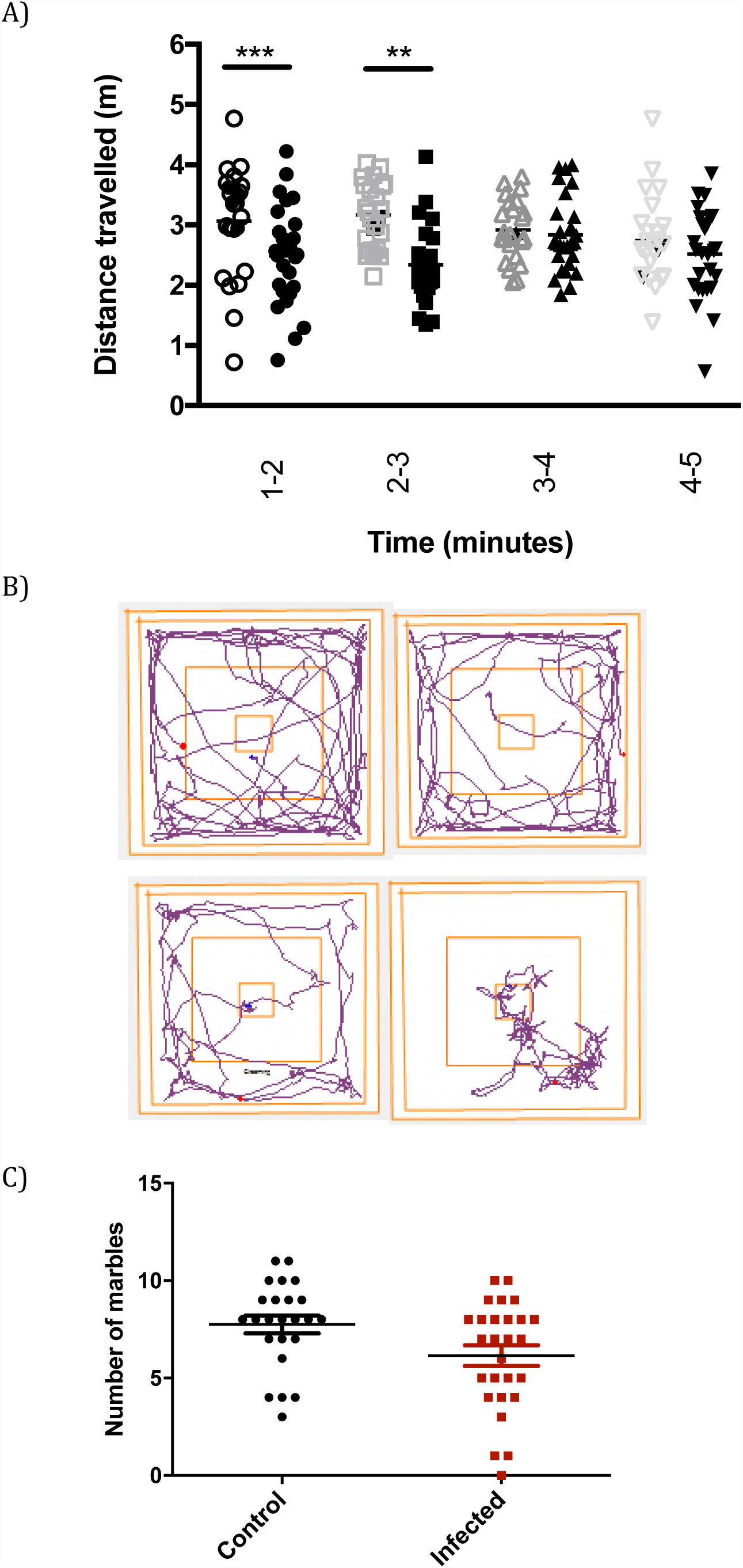
Locomotion and anxiety-related behaviour are altered in infected animals. A) Ambulation of uninfected (grey) and infected mice (black) in the open field at single minute timepoints with the mean. **, p=0.0015; ***, p=0.000097, student t-test. B) Tracking in the open field for representative uninfected (top) and infected (bottom) mice from 0-180 seconds of the trial. C) Plot of the number of marbles buried during marble burying trials for 30 minutes with uninfected (black circles) and infected mice (red) (p=0.028, t-test, n=51).

Disruption of noradrenergic signalling has also been associated with anxiety changes, although anxiety is a complex cognitive process with the contribution of multiple neurotransmitters. Marble burying is an anxiety-related behavior in mice, where repetitive digging response is a defensive trait (44). In our trials, infected mice buried a reduced number of marbles compared with uninfected mice (Figure 5C; p= 0.028). There was very little association between marble burying and DBH mRNA level (Figure S4, Correlation coefficient = −0.18). The minor change observed here fits with conflicting observations of changes in anxiety-related behavior with *T. gondii* infection found in the literature; with reduced fear observed in open spaces in the elevated plus maze reported, while others found no effect in the open field (2, 39, 45, 46). It has also been suggested that *T. gondii* may damage hippocampal function, since hippocampal neurons and glial cells may be infected, so differences in marble burying could reflect changes in hippocampal function (47).

### Effect of Sex on Altered Norepinephrine Regulation with Infection

An anomaly that was noted in testing was a large variation in DBH mRNA levels in the brains of female animals. The large range of DBH levels would mask any effect by infection. Indeed, infected females did not exhibit a measurably lower level of DBH (Figure 6A, p=0.45) with infected females possessing higher and lower DBH mRNA levels than vehicle controls (Table 1). We investigated the reasons for this difference. DBH gene expression is regulated by estrogen, with the estrogen receptor binding to ER-response elements (ERE) at the 5’ flanking region of the DBH gene and activating transcription (48, 49). Estrogen, estrogen receptor and DBH mRNA levels fluctuate during the estrous cycle (50). Hence, we measured the levels of estrogen receptor 1 (ESR1) mRNA in the brains of the female rats used in this study.

**Figure 6:**
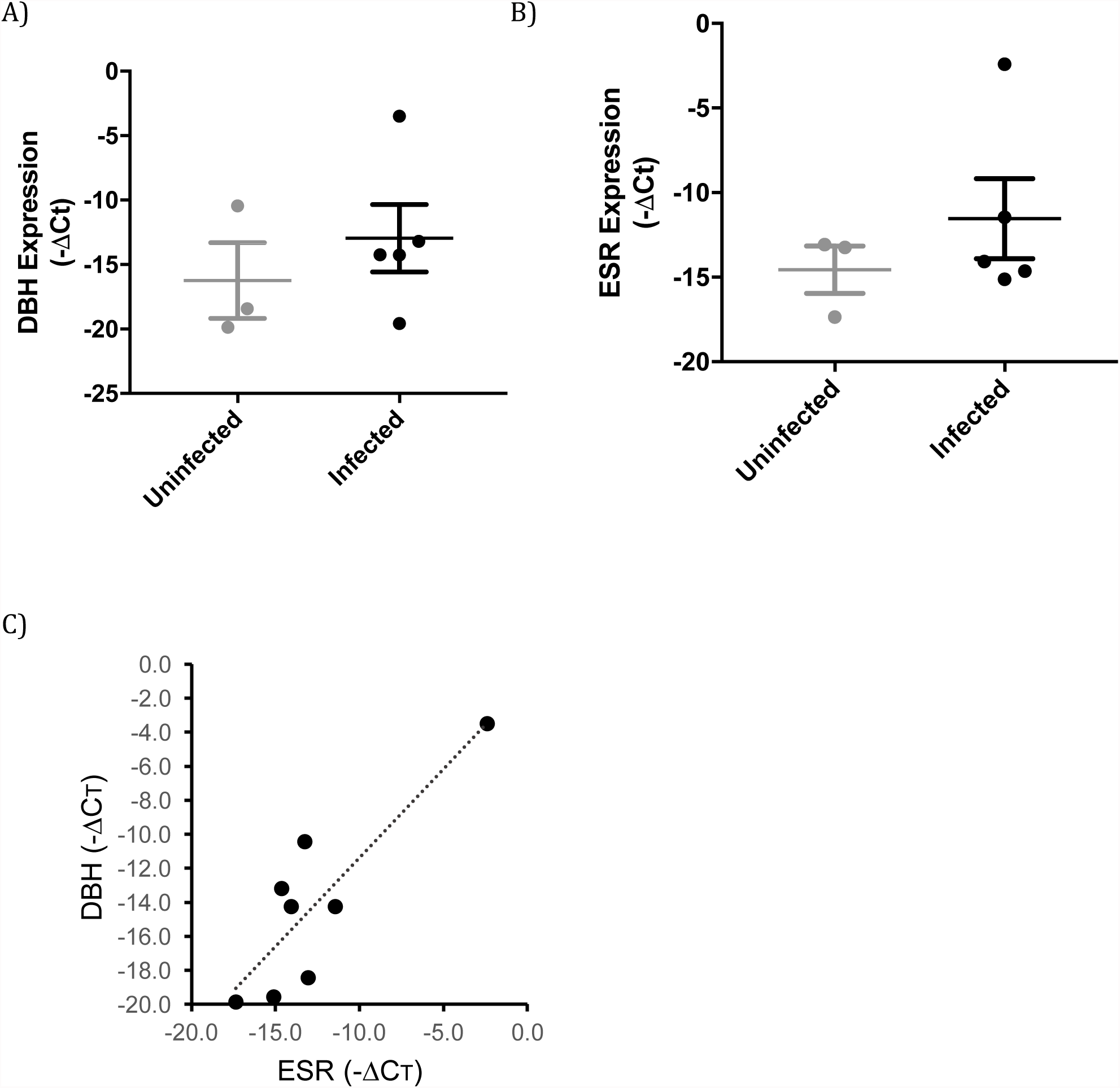
Dopamine ß-hydroxylase expression was not suppressed in infected female rats. A) A plot of DBH mRNA in the brains of uninfected (grey) and chronically infected (black) female rats is plotted; ±SEM; n=8; student t test p=0.45. B) The expression of the estradiol receptor 1 (ESR1) gene in brains of uninfected (grey) and chronically infected (black) female rats shown graphically; ±SEM; n=8; student t test p=0.40. C) The level of DBH plotted versus the level of ESR1 gene expression in brain sections of rats in this study (Pearson’s correlation coefficient = 0.86).

A range of ESR1 levels were observed in the brains of the female rats, indicative of differences in their estrous cycle (Table 1). Expression of ESR1 was not altered by infection (Figure 6B, p=0.40). ESR1 mRNA levels, however, strongly correlated with DBH mRNA (Figure 6C), with a correlation coefficient of 0.86 (p=0.0064), as expected (50). Together, the findings show that DBH expression correlated with ESR1 expression but not infection in females. These findings provide a biological basis for previously observed sex-specific differences in the effect of *T. gondii* infection on mouse behavior and estrous-dependence of aversive behaviors in female rats (51, 52).

### Dopamine β-hydroxylase expression in cytomegalovirus infected human neuronal cells

To test whether DBH down-regulation is a general response to CNS infection or whether it is specific, changes in DBH gene expression in human neuronal cells infected with human cytomegalovirus (HCMV) were measured. DBH mRNA levels were not significantly changed over a time course of HCMV infection in BE(2)-M17 cells (p>0.13), with a trend for increased expression at 24 hours (Figure 7A). At this point, HCMV is entering the late stages of viral replication (as indicated by the immediate-early UL123 gene expression in Figure 7B) and yet the data clearly show HCMV infection does not decrease DBH expression. In comparison, DBH gene expression was down-regulated (relative to the marker) in the same cells infected with *T. gondii*, decreasing over the time course of the experiment (Figure 7C) with a small increase in the number of *T. gondii* (Figure 7D). Hence, DBH down-regulation is specific for *T. gondii* infection.

**Figure 7:**
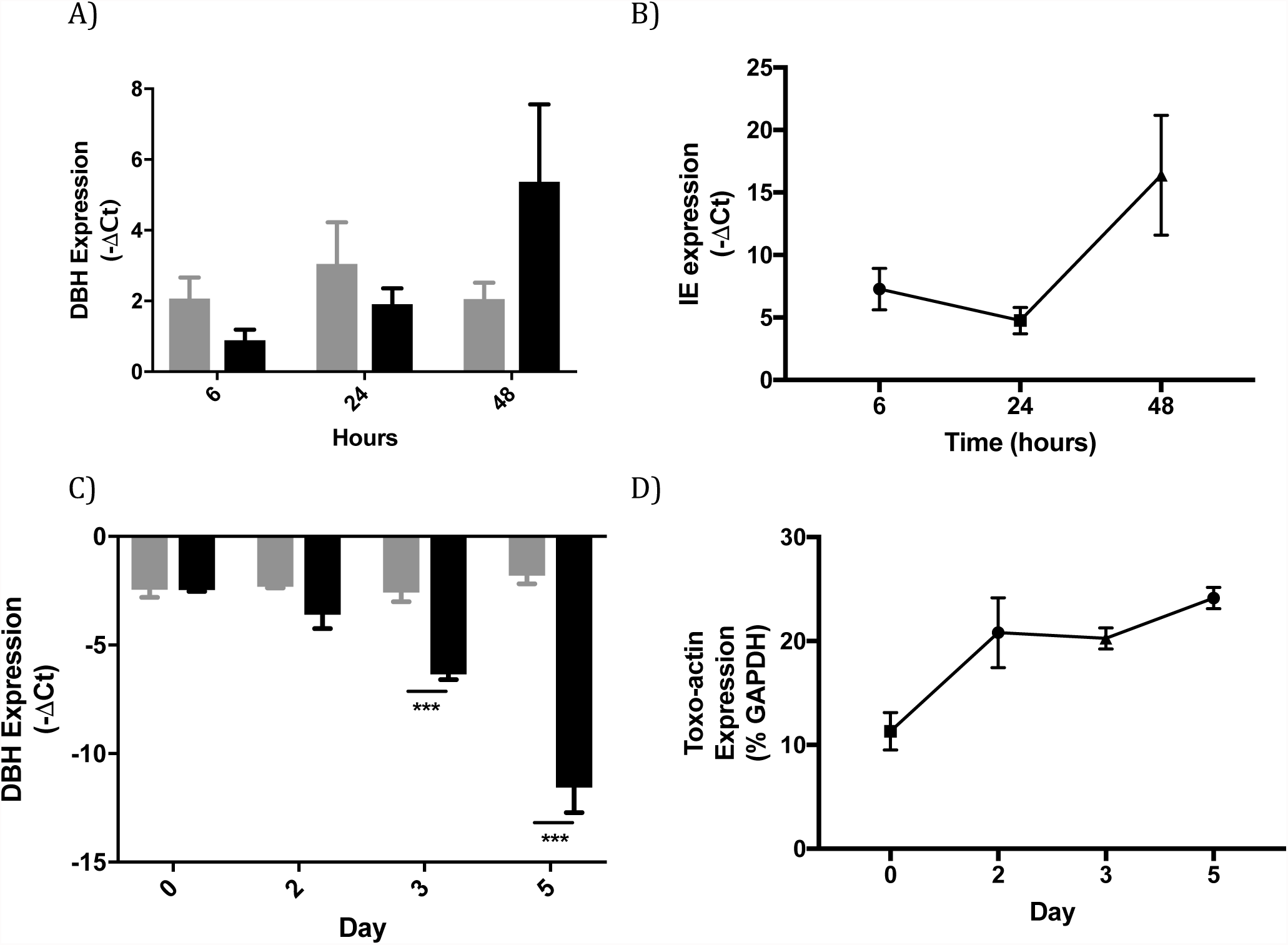
Dopamine ß-hydroxylase suppression is pathogen-specific. A) Plot of DBH gene expression over a time course of 48 hours. Uninfected (grey) and human cytomegalovirus (CMV) infected (black) human BE(2)-M17 neuronal cell line, shown as a percentage of the housekeeping gene GAPDH. Multiplicity of infection is 1; n=2. B) Accumulation of HCMV UL123 immediate-early (IE) as percent gene expression (normalized to GAPDH) over a time course. C) Plot shows DBH expression over a similar time course for uninfected (grey) and *T. gondii* infected (black) human neuronal cells, as a percentage of the housekeeping gene GAPDH. Multiplicity of infection is 1; ***, p=0.0015 and 0.0012, respectively, Student’s t test; n=3; error bars indicate SEM. D) The intensity of *T. gondii* infection over the time course based on levels of *T. gondii* actin plotted as a percentage of host GAPDH.

## Discussion

A decrease in the neurotransmitter NE was observed in *T. gondii*-infected brains, and, for the first time, the down-regulation of expression of the DBH gene, that encodes the key enzyme in NE synthesis, was discovered as the mechanism responsible. This study examined changes in gene expression throughout the genome to identify neuronal changes with infection rather than focus on specific neural genes. Levels of DBH gene expression were highly correlated, inversely, with infection intensity. Prior studies of neurotransmitters in neurons during infection found elevated levels of metabolites of neurotransmitters and alterations in neurotransmission but did not identify the mechanisms responsible for altered neurotransmitter levels (8, 11–13, 18). Further, DBH expression was down-regulated >30-fold with chronic infection (Figure 3). In other studies, GABA and glutamate metabolism in the CNS of chronically-infected animals were altered. A change in the distribution of the GABA-associated protein GAD67 was found in neurons of infected animals but GABA levels were not measured (53). Elevated levels of CNS glutamate were also found at 35 and 42 days post-infection in mice and associated with a 50% reduction in GLT-1 expression in astrocytes (54). Hence, multiple mechanisms including immunological and direct changes in neurons are responsible for neurophysiological changes in the CNS with *T. gondii* infection.

In this study, the changes in catecholamine regulation observed provide a mechanism that can resolve the diverse observations of CNS catecholamines with infection. The large down-regulation in DBH gene expression in the *T. gondii*-infected brain and in catecholaminergic neural cells observed will disrupt a key step in catecholamine metabolism (Figure 2, 3). This down-regulation will decrease metabolism of DA into NE, resulting in lower NE levels and elevated DA levels. Indeed, the DBH suppression observed corresponds with the decreased NE and concurrent increase in DA in infected PC12 cells, where no changes have been found in amounts of other enzymes in the biosynthetic pathway (Figure 2, (8)). Down-regulation of DBH expression also provides an explanation for the observed decreases in NE in infected brains, but without a significant increase in DA in brain tissue (Figure 1) since only neurons expressing DBH will be affected. This is unsurprising given the proportions of dopaminergic and noradrenergic neurons in the brain. This, combined with the more severe pathology of *T. gondii* infection in mice with dysfunctional neurons, may also explain other studies that did not detect changes in brain DA levels with infection (11, 15, 28, 32, 55). Down-regulation of DBH gene expression was specific for *T. gondii* infection and not due to apoptotic or necrotic responses (Figure 3C) and was not observed with infection by the neurotropic pathogen CMV (Figure 7). Suppression of DBH and NE was only observed in males, while expression of the estrogen-regulated DBH gene correlated with ESR1 levels in females (Figure 6).

The down-regulation of DBH found in this study can account for the increased DA observed in infected PC12 cells observed in earlier studies (8, 18). In those studies, the amount of DA increased with infection while levels of the enzymes in synthesis, tyrosine hydroxylase and dopa decarboxylase were unchanged, although dopa decarboxylase could be detected in the parasitophorous vacuole. *T. gondii* contain two paralogous genes that encode an aromatic amino acid hydroxlase (TgAAAH), with tyrosine and phenylalanine hydroxylase activities, that is secreted from the parasites into the parasitophorous vacuole (21). Both paralogs were found to be expressed in bradyzoites, whereas only TgAAAH1 was expressed in tachyzoites. The gene products have been found to be involved in oocyst development as proposed in their original discovery (21, 22). The effects of disruption of one of the two paralogs on catecholamine neurotransmission remain inconclusive; hence, collaborative experiments using the recently developed double knockout mutants lacking both genes are ongoing (22).

Noradrenergic neurons are principally located in the locus coeruleus (LC) in the brain and project to the thalamus, hippocampus and the frontal and entorhinal cortices, as well as, to a minor extent, most other brain regions (56). *T. gondii* cysts have been observed in these brain regions (57, 58). LC terminals can release both NE and DA, and, recently, efferent noradrenergic neurons originating in the LC were found to release DA in the dorsal hippocampus (59, 60). In this study, DBH gene down-regulation with chronic infection was observed in the PFC and LC/pons regions (Figure 3B). With the DBH suppression observed in this study, noradrenergic neurons may have increased DA released relative to NE.

With decreased NE in the brain with infection, changes were observed in noradrenaline-related behaviors. Infected mice exhibited down-regulation of DBH in the brain, associated with decreased arousal and increased social interactions, with DBH level in the infected mice correlating with behaviors (Figures 6,7). Anxiety was also reduced in the marble burying task with infection. Chronic *T. gondii* infection has also been found to impair long-term fear memory, a process that NE enhances (11, 61). Although one could attempt to reverse the parasite-induced effects on noradrenaline-related behaviors with noradrenergic inhibitors, antipsychotic drugs have antiparasitic effects (28, 62, 63), and L-threo-3,4-dihydroxyphenylserine cannot be used because the required dopa decarboxylase for activation is altered by *T. gondii* infection (8, 64).

There is a link between NE levels, *T. gondii* infection and movement and coordination of the host. Both *Dbh*−/− knockout in mice and noradrenergic neuron loss in the LC (in rats) lead to motor impairments and development of dyskinesia (65, 66). Further, mice lacking NE are susceptible to seizures (67, 68). Chronic infection with *T. gondii* in mice has also been associated with coordination difficulties (69), and loss of coordination is a common symptom of human toxoplasmosis. Severe toxoplasmosis can cause seizures, with documented cases of patients exhibiting Parkinsonian traits such as bradykinesia (70, 71). Effects of altered GABA metabolism with *T. gondii* infection (observed in an earlier study) in promoting seizures would be compounded by a lack of anticonvulsant effect promulgated by NE (53).

Although DBH gene expression strongly correlated with the intensity of infection (Figure 4), the low number of encysted neurons and lack of apparent tropism (data not shown) is difficult to reconcile with the large decrease in DBH expression. The numbers of cysts found in this study were similar to a previous study of 105 *T. gondii*-infected rats (72). The large effect with relatively low numbers of cysts is similar to observed global changes in GAD67 (glutamic acid decarboxylase) distribution in the brains of *T. gondii*-infected mice (53). These changes could be mediated by injection of parasite proteins into cells without infecting the cells, as has been observed with neurons in infected mice (7, 73). This will be the subject of future studies.

Infection of the CNS can influence brain neurophysiology, as found here with NE levels. *T. gondii* infection was discovered to down-regulate DBH gene expression, tightly correlating with infection intensity. This can result in suppressing NE while elevating DA in the same neurons. Further studies need to define the consequential effects on neurological signalling of these alterations as they will depend upon the location of the noradrenergic neurons and dopamine receptors. The mechanisms whereby the parasite down-regulates DBH expression need clarification. This may be via a parasite mechanism similar to *T. gondii* ROP18 altering JAK/STAT signaling pathways or via the regulation of vasopressin receptor by epigenetic changes (74, 75). The neurophysiological changes observed may provide insights into the mechanisms responsible for behavioral effects of *T. gondii* infection (76).

## Materials and Methods

### Ethics

All procedures were approved by the University of Leeds Animal Ethical and Welfare Review Board and performed under United Kingdom Home Office Project and Personal Licences in accordance with the Animals (Scientific Procedures) Act, 1986. Rat brain sections were from infections conducted at the School of Public Health, Imperial College London (ICL) and procedures were approved by the ICL Animal Care and Use Committee and following the same Home Office, HSE, regulations and guidelines. Considerations of replacement, reduction, and refinement were taken in the use of animals for research.

### Rodent and rodent infections

The (BALB/cAnNCrl × C57BL/6NCrl)F_1_ mice used in this study were bred by crossing C57BL/6NCrl males to BALB/cAnNCrl females (Charles River Laboratories). The C57BL/6 inbred strain has been used as the genetic background in prior behavioral studies of *Dbh*−/− knockout mice, while the BALC/c inbred strain possesses genetic resistance to control *T. gondii* brain infection and develops a latent chronic infection (77). In pilot studies, purebred C57BL/6NCrl mice infected with *T. gondii* showed severe toxoplasmic encephalitis.

Mice were housed five of the same sex per cage, with *ad libitum* access to food pellets and water. Mice were checked for health changes daily and their weight was measured weekly. Any mouse showing severe illness or significant weight loss (25%) was promptly culled. Mice were grouped according to treatment. Mice were infected by intraperitoneal (IP) injection with *T. gondii* type II strain Prugniaud in sterile phosphate-buffered saline (PBS) at 6–14 weeks of age. Infection was monitored by the direct agglutination test (BioMérieux) to detect *Toxoplasma* antibodies, following the manufacturer’s instructions, in sera from collected blood samples. Brains were harvested from euthanized animals and snap frozen. Cryosectioned slices were used for RNA isolation as described for rats below.

Rat samples were from Lister Hooded rats (Harlan UK Ltd), males and females housed separately and provided food and water *ad libitum*, that were infected at approximately 3 months of age via IP injection of 1 × 10^6^ tachyzoites in sterile PBS. Uninfected control rats were IP injected with sterile PBS and sacrificed 5-6 months post-infection, with brains quick-frozen for cryosectioning. Sagittal slices were processed for RNA by dissolution with Trizol™ (Thermo Fisher) for processing following manufacturer’s instructions.

### Growth of pathogens and cultured cells

The *T. gondii* Prugniaud strain was maintained in human foreskin fibroblast cell line Hs27 (ECACC 94041901), as previously described (21). Rat adrenal phaeochromocytoma (PC-12) cells (kind gift from C. Peers; ECACC 88022401) were maintained in RPMI (Invitrogen, Paisley, UK), supplemented with 10% horse serum (Invitrogen), 5% fetal bovine serum (FBS; Invitrogen), and 100 units/ml penicillin/streptomycin (Sigma, Poole, UK). PC-12 cells were passaged by triturating, centrifuging 800 rpm for 10 min in a table top centrifuge, resuspending in fresh media and incubating at 37°C in an atmosphere of 5% CO_2_.

For the induction of parasite conversion to bradyzoite forms, free released tachyzoites were incubated at 37°C in RPMI supplemented with 1% FBS (pH 8.2) for 16-18 hours (hr) in ambient air then diluted with DMEM (Invitrogen), isolated by centrifugation, and suspended in RPMI (pH 7.4) containing horse serum, FBS and penicillin/streptomycin, as previously described (18).

For HCMV studies, cells were infected with wild type Merlin HCMV strain for 1 hour then washed and incubated with fresh media. RNA was harvested at the times shown. Cells were confirmed permissive for HCMV by IE antigen staining, which demonstrated similar susceptibility for infection as the neuronal cell line U-373, an established permissive HCMV cell line.

### RNA sequencing and data analysis

PC-12 cells were cultured in poly-D-lysine-coated 6-well plates (Sigma). Following 24 hours of incubation, 6 × 10^4^ cells were changed to medium with 1% horse serum, 0.5% FBS. After a further 24 hr, 100 ng/ml of Nerve Growth Factor (NGF; Sigma) was added. The addition of NGF was repeated once every 24 hr throughout the length of the experiment. Control experiments found no effect of NGF on growth or bradyzoite conversion of *T. gondii* (data not shown). After 72 hr from the initial addition of NGF, dendritic extensions were visible from differentiated cells. At this point, induced Prugniaud tachyzoites were transferred to each well, maintaining a parasite density of 2.5 × 10^4^ cells/ml. Cells were harvested immediately following infection (day 0) and after three and six days of infection for RNA extraction. The cultures were monitored daily by light microscopy. At day 6 of infection, the parasitaemia level was 60-70%, with little observable cell lysis (data not shown).

Cells were detached from the surfaces by manual removal with a scraper and several parallel biological repeats were pooled. The suspended cells were pelleted by centrifugation at 800xg for 10 minutes and lysed with TRI Reagent solution (Invitrogen) followed by centrifugation at 12,000xg for 10 minutes at 4°C. RNA was purified following manufacturer’s instructions. RNA samples were stored at −80°C.

mRNA was enriched using a Poly(A)Purist™ MAG Kit (Ambion) followed by further enrichment using RiboMinus™ (Ambion), following manufacturer’s instructions. Following quality control analysis using a Bioanalyzer (Agilent), cDNA libraries were prepared from RNA using the Epicentre ScriptSeq v2 RNA-Seq Library Preparation Kit and sequenced using the Illumina Hiseq 2000 at the University of Liverpool Centre for Genomic Research. Two libraries for each pool of biological repeats were sequenced. RNA sequencing generated 353m paired-end reads, with a total of 26,405 *Rattus norvegicus* genes identified.

The Illumina reads from the RNA sequencing were separately mapped to *Rattus norvegicus* and *Toxoplasma gondii* reference genomes using Tophat 2.0.8b (78). Differential expression analyses were performed using edgeR package version 3.0.4 (79) for the reads aligned to the rat genome. A gene was considered as differentially expressed (DE) if the fold change was greater than two (−1 > log2(fold change) > 1) and the FDR < 0.01. The resultant 488 genes form a reliable set of DE genes that exhibit down- or up-regulation (Table S1). The enriched GO (Biological Process) and KEGG pathway terms for up- and down-regulated gene sets were computed using DAVID and are tabulated in Table S1 (80).

### Reverse transcriptase PCR and quantitative PCR

For RT-qPCR assays, cultures of 2.5 × 10^4^ PC-12 cells in multiwell plates were infected with induced *T. gondii* tachyzoites. PC-12 cells were infected with multiplicities of infection (MOI); after five days, cells were recovered by centrifugation and the cell pellet frozen (−80°C) for RNA extraction and HPLC-ED analysis.

RNA was purified using Direct-zol™ (Zymo) and reverse transcribed to cDNA using Maxima First Strand cDNA Synthesis Kit (Thermo Fisher), following manufacturer’s instructions. RT-qPCR was performed on RNA, as described previously, using SYBR^®^ Green Real-Time PCR Master Mix (Thermo Fisher) using rat GAPDH primers (Qiagen), DBH primers 5’-CCACAATCCGGAATATA-3’ and 5’-GATGCCTGCCTCATTGGG-3’, and ESR primers 5’-CTACGCTGTACGCGACAC-3’ and 5’-CCATTCTGGCGTCGATTG-3’.

### HPLC for monoamines

The catecholamines DA and NE were measured by HPLC-ED, adapting a previously published method (18). Briefly, cultures were harvested by scraping cells, recovered by centrifugation, and an aliquot taken for cell counting and normalization. The remaining cells were recovered again and resuspended in 350 µL of perchloric acid, followed by sonication. The mixture was centrifuged at 14,000 rpm for 15 minutes at 4°C to remove particulates, and an aliquot was taken for HPLC analysis. NE was detected at 4.5 minutes and DA at 8 minutes (flow rate 0.4ml/min) by HPLC-ED on a Dionex UltiMate 3000 system (Thermo Fisher).

### Mouse Behavioral Testing

After establishment of chronic infection (4-5 weeks), mice were tested in a battery of behavioral tests in the following order, with an interval of 2 days between each test: open field > marble burying > social approach. Prior to testing, mice were habituated to handling for 5 minutes per day for 7 days. Ethanol (70%) was used to clean the arena between mice. The arena was left to dry for 3-4 minutes before commencing the next subject.

### Open Field Test

The internal open field arena had a diameter of 40 × 40 cm with a semi-transparent Perspex wall. The arena floor was white plastic. To prevent the mice from seeing the surrounding room, a cylinder of white card was placed around the arena 30 cm away from its walls. The ambulation of the mice was recorded using a webcam that was placed on a tripod above the arena.

Mice were individually placed at the centre of the arena facing the same wall. Readings began after the initial 60 seconds because of disturbances involved in the experimenter removing mice from their cages, placing them in the open field and withdrawing to a computer to manually start the recording. Distance travelled was recorded for 15 minutes without interruptions or intervals. using AnyMaze tracking software (Stoelting Co.).

### Social Approach

Sociability was assessed using a three-chambered arena (60 × 40 cm) that had two openings (7 × 8 cm) to allow the mouse access to the left and right chambers from the central chamber (each chamber measured 40 × 20 cm). The test involved using two unfamiliar mice that had been habituated to stainless steel cylinders (10 cm W × 10.5 cm H) prior to the test. The cylinders were made of vertical metal bars separated by 9 mm, which allowed air exchange and increased the possibility of contact between the test and stranger mice.

Following a previously published protocol (37), a test mouse was placed into the central chamber of the three-chambered arena. The ‘habituation’ stage was carried out for 15 minutes; at the end of this time, the test mouse was moved to the central chamber and the openings to the side chambers were blocked by guillotine doors. A cylinder was placed in both the right and the left chamber. A stranger mouse (‘stranger 1’, a young male C57BL/6NCrl) was placed in the cylinder in either the left or right chamber (balanced between treatment groups). Following this, the doors were removed and ‘phase 1’ was initiated, lasting 10 minutes.

Social approach was scored when the test mouse’s nose poked through the bars of either the cylinder containing stranger 1 or the empty cylinder. At the end of phase 1, the test mouse was placed in the central chamber and the doors were shut. Then, a new unfamiliar mouse (‘stranger 2’) was placed in the formerly empty cylinder. At this point, phase 2 was initiated, again lasting for 10 minutes. Social approach was scored when the test mouse’s nose poked through the bars of either the cylinder containing stranger 1 or the cylinder containing stranger 2. The cylinders and floor were then wiped clean with 70% ethanol. The experimenter wore nitrile gloves throughout the procedure.

### Marble Burying

In a large cage (rat cage), 12 glass marbles were placed in a consistent grid pattern on wood-chip bedding that was lightly tamped down to make a flat, even surface. The mouse was placed in the cage and left for 30 minutes. The number of marbles buried up to two-thirds of their depth was counted after 30 minutes.

### Statistical Analysis

GraphPad Prism (Version 7) was used for statistical analyses. All data are plotted mean ± SEM.

## Authors’ contributions

The main manuscript text was written by I.A., E.T. and G.M., with input from all authors. I.A. and E.T. contributed equally to this study. Experiments were performed and figures and tables prepared by I.A., E.T., M.A., G.B. and M.S.V. I.A., G.M. and J.W. contributed to the conceptualization and experimental planning. M.R. is supported by MRC Fellowship G:0900466. The Stanley Medical Research Institute supported early components of this study.

